# ApoC-III helical structure determines its ability to bind plasma lipoproteins and inhibit Lipoprotein Lipase-mediated triglyceride lipolysis

**DOI:** 10.1101/2020.07.02.183178

**Authors:** Cecilia Vitali, Sylvia Stankov, Sumeet A. Khetarpal, John Millar, Leland Mayne, S. Walter Englander, Nicholas J. Hand, Sissel Lund-Katz, Michael C. Phillips, Daniel J. Rader

## Abstract

In humans, apolipoprotein C-III (apoC-III) plasma levels have been associated with increased risk of cardiovascular disease. This association is in part explained by the effects of apoC-III on triglyceride (TG) metabolism; apoC-III raises plasma TG by increasing very low density lipoprotein (VLDL) secretion, inhibiting lipoprotein lipase (LPL)-mediated TG lipolysis, and impairing the removal of triglyceride-rich lipoprotein (TRL) remnants from the circulation. In this study, we explored the structure-function relationship the interaction of apoC-III with plasma lipoproteins and its ultimate impact on LPL activity. The structural and functional properties of wild-type (WT) apoC-III were compared with two missense variants previously associated with lower (A23T) and higher (Q38K) plasma TG. ApoC-III in the lipid-free state is unstructured but its helix content and stability increases when bound to lipid. Lipid-bound apoC-III contains two alpha helices spanning residues amino acids 11 - 38 (helix 1) and 44 – 64 (helix 2). Investigation of the structural and functional consequences of the A23T and Q38K variants showed that these amino acid substitutions within helix 1 do not significantly alter the stability of the helical structure but affect its hydrophilic-lipophilic properties. The A23T substitution impairs lipoprotein binding capacity, reduces LPL inhibition, and ultimately leads to lower plasma TG levels. Conversely, the Q to K substitution at position 38 enhances the lipid affinity of helix 1, increases TRL binding capacity and LPL inhibition, and is associated with hypertriglyceridemia. This study indicates that structural modifications that perturb the hydrophilic/lipophilic properties of the alpha helices can modulate the hypertriglyceridemic effects of apoC-III.

## Introduction

High levels of plasma triglyceride (TG) are associated with greater incidence of cardiovascular disease (CVD)^1, 2^ and triglyceride-rich lipoproteins (TRL) are known to be a causal risk factor for CVD ^3, 4^. Among the factors affecting TRL metabolism is apoC-III, an 8.8 KDa exchangeable apolipoprotein primarily secreted by the liver^5^. ApoC-III exchanges between TRL and HDL, and higher concentrations of the protein lead to increased plasma TG levels^6^. Consistent with a critical role for apoC-III, genome-wide association studies have shown that loss-of-function (LOF) variants of this protein are associated with reduced plasma TG levels ^7–9^. These observations suggest that inhibition of apoC-III function should be of therapeutic benefit and there is much interest in developing suitable approaches^10^. In particular, reduction of apoC-III expression using an antisense oligonucleotide ^11, 12^ and direct protein targeting with a monoclonal antibody ^13^ show promise.

There is a long history of studies aimed at elucidating the mechanisms underlying the modulation of plasma TG levels by apoC-III. An early investigation of hypertriglyceridemic mice expressing human apoC-III showed that there is decreased tissue uptake of TRL particles from the circulation, most likely due to increased apoC-III and decreased apoE on VLDL particles^14^. Consistent with this concept, studies with antisense oligonucleotides targeting apoC-III lowered plasma TG in both mice and humans through a mechanism dependent upon the family of LDL receptors^15, 16^. Besides its effects on receptor-mediated clearance of TRL remnant particles, apoC-III in normal human subjects can inhibit the catabolism of TRL by lipoprotein lipase (LPL) thereby raising plasma TG levels^17, 18^. Further insight into the functionality of apoC-III has been gained by comparing the behavior of mutants to wild-type (WT). Thus, studies of the LOF A23T apoC-III variant which exhibits impaired lipid binding ^19^ confirm that hydrolysis of TG in TRL is enhanced when less apoC-III is present ^13^. ApoC-III has also been reported to affect the secretion of VLDL-TG from liver ^20, 21^. In this regard, the A23T LOF variant has provided conflicting results, showing an association with impaired VLDL-TG secretion when expressed in cells ^19^ but no appreciable effect when expressed *in vivo* ^13^ whereas the Q38K gain-of-function (GOF) variant has been shown to increase VLDL assembly and secretion ^22, 23^. Overall, it is apparent that apoC-III can modulate plasma TG levels by influencing both the number of TRL particles in circulation (altered secretion and/or clearance) and their composition (altered TG hydrolysis by LPL). The relative contributions of these processes to the overall control of plasma TG levels is still unclear and apparently model dependent. In this study, we aimed at providing mechanistic insight into the relationship between apoC-III structure, its lipid-binding properties, and the inhibition of LPL-mediated TG lipolysis. Results indicate that binding of apoC-III to TRL particles is crucial for inhibition of LPL activity and that this process is accompanied by helix formation within the protein. Furthermore, the study of the structural and functional properties of the A23T and Q38K variants showed that amino acid substitutions altering the hydrophilic-lipophilic properties of the amphipathic α-helices can significantly impact the ability of ApoC-III to bind to TRL, even in absence of major modifications of the stability of the α-helical structures. Structural modifications that reduce the lipid binding ability of apoC-III have the potential to lower the hypertriglyceridemic effects of the protein.

## Materials and Methods Mice

Apoc3-knockout (KO) and WT mice on a C57BL/6J background were obtained from the Jackson Laboratory and bred at the University of Pennsylvania. The stock numbers for these strains are 002057 and 000664, respectively. Mice were fed *ad libitum* with a standard chow diet and were maintained on a 12-h light cycle. All procedures were conducted according to IACUC-approved protocols.

### Recombinant proteins

Recombinant apoC-III proteins were purchased from DNA-Protein Technologies LLC in Oklahoma City (http://www.DNA-Protein.com); these proteins were prepared as follows.

Human apoC-III cDNA cloned into the pET32a(+) vector from Novagen was used for expression of wild-type (WT) apoC-III with a His-tagged thioredoxin fusion protein at the amino terminal^24, 25^. A tobacco etch virus protease site engineered into the expression vector allowed for removal of the thioredoxin fusion protein without any extra amino acids being added to the native human apoC-III sequence^26, 27^. cDNA inserts encoding the human apoC-III variants A23T and Q38K were engineered from the above pET32a(+) plasmid using the QuikChange Site-Directed Mutagenesis Kit (Stratagene, CA). The human apoC-III variants encoded by these cDNA inserts were expressed in *E. coli* strain BL21-DE3 and purified (>95% purity) according to previously published procedures^24, 25^. The above apoC-III proteins were also purchased from Thermo Fisher Scientific at the same level of purity as peptides prepared by solid phase synthesis. Unilamellar vesicles of dimyristoyl phosphatidylcholine (DMPC) (Avanti Polar Lipids, Alabaster AL) at 10 mg/ml in aqueous buffer were prepared by sonication using established procedures^28^.

### Intralipid clearance

The ^3^H-Intralipid substrate used for clearance studies was prepared as follows. [^3^H]triolein in toluene (PerkinElmer) was evaporated under a nitrogen gas. Intralipid 20% (Fresenius Kabi) was added to the dried ^3^H-triolein and allowed to equilibrate overnight at 4°C with gentle shaking.

The mixture was then incubated for 1h at 37°C with either hApoC-III dissolved in PBS (13.5 mg/dL in final mixture) or equal volume of PBS (negative control). 11-14 week old male mice were intravenously injected with a dose of 100uL of such substrate, containing approximatively 8.4-9.7×10^6^ CPM. Plasma was collected at various time points after ^3^H-TRL administration (1-122 min). ^3^H activity of plasma from each time point was measured by scintillation counting. The relative ^3^H activity was calculated by normalizing the activity from each time point by that at 1 min.

### Hydrogen Exchange and Mass Spectrometry

Hydrogen-deuterium exchange (HX) coupled with fragment-separation^29^ and mass spectrometry (MS) analysis^30–33^ was used to compare the secondary structure of wild-type (WT) and variant forms of human apoC-III free in aqueous solution and bound to DMPC vesicles. The fragment-separation system consisted of a box maintained at 0°C with an immobilized pepsin column, a C18 trap and an analytical C18 column connected in the order listed; for more details on the fragment separation system see reference^32^. The outlet from the C18 analytical column was connected in line with an LTQ Orbitrap XL mass spectrometer (Thermo Fisher Scientific) for analyzing the peptide mass (MS analysis) and peptide sequence (MS^2^ analysis). HX time-point samples were studied at pD 5.8 (pH 5.4) and 5°C. All solutions were equilibrated at the desired temperature and HX was initiated by diluting the stock apoC-III in H_2_O buffer (50mM (NH_4_)_3_PO_4_, pH 5.4) 10-fold into 100% D_2_O buffer (50mM (NH_4_)_3_PO_4_, pD 5.8) at 5°C to yield a final apoC-III concentration of 2.5-5 μM (20 – 40 μg/ml). Low temperature and acidic conditions were used to slow the apoC-III HX rate so that D incorporation at short times could be measured. The exchange reaction was quenched at specific time-points by adding a pre-calibrated amount of 99% formic acid on ice to lower the pD to 2.3. A 50μl aliquot of the quenched apoC-III HX sample was immediately injected into the fragment separation system. Between 0-3 min after injection the proteins were digested in the pepsin column and washed in the C18 trap column. The peptides were then roughly separated on an analytical C18 column (flow rate 7μl/min) with a water-acetonitrile gradient (12-50% acetonitrile) (pH 2.3, 0°C) over 3-15 min. Eluant was injected by electrospray into the spectrometer for mass analysis. Samples containing 100% D to calibrate back-exchange were run and a MS^2^ run on a 100% H sample was performed to calibrate the peptide retention times.

Mass spectra from the HX experiments were analyzed using in-house software ExMS^34^. The number of D incorporated at each time-point was calculated from the centroid values. This number for each peptide fragment was corrected for back-exchange and used to create a measured HX rate curve (*k*_obs_) for that fragment. The back exchange was calculated as fraction D retention = (((100% D centroid - 100% H centroid) × charge state)/number of exchangeable amides in the peptide). The number of deuterons at each HX time point was calculated from the centroid value as, number of D = (((centroid of time-point - centroid 100%H) × charge state)/fraction D retention). The measured HX rate curve was compared to a predicted rate (intrinsic or reference rate, *k*_ref_) curve^35–37^ assuming the fragment to be in a dynamically disordered random coil state so that the amide hydrogens are not protected against exchange with water. The intrinsic rate and observed data were fit to stretched exponentials (a stretching factor is included because the hydrogens in any given amino acid sequence exchange over a range of rates determined by nearest neighbor effects)^30^ and a protection factor (Pf) was calculated as a ratio of reference rate/observed rate (Pf = *k*_ref_/*k*_obs_). For more details on data analysis see references^30, 38^. Any apoC-III fragment with Pf <10 is considered to be unprotected (disordered) and any fragment with Pf > 10 is considered to be protected^30, 39, 40^. For peptide fragments with an average Pf > 100, the amide hydrogens are involved in hydrogen bonded secondary structure such as α-helix that is relatively stable. Assignment of structure to peptide fragments with Pf in the range 10 – 100 is ambiguous; such values can arise from either random coil structure that is not completely dynamically disordered or very unstable α-helices that are unfolded for a significant fraction of the time, as shown in other protein systems^41, 42^. Pf was used to calculate the free-energy of HX (∆*G*_HX_= - RT ln (*k*_obs_/*k*_ref_) = RT ln Pf) which corresponds to the free energy of concerted helix unfolding and is therefore a measure of helix stability.

### LPL activity assay

In vitro LPL activity assay was determined using a TRL-like lipid emulsion as substrate, according to the protocol described by McCoy et al. with minor modifications^43^. Briefly, the concentrated substrate was prepared by mixing 300 mg of triolein (Sigma-Aldrich), 18 mg of egg-phosphatidylcholine (Sigma-Aldrich) and 1.12 mCi of ^3^H-triolein (PerkinElmer) were dried under a nitrogen gas. Lipids were combined with 5 ml of glycerol (Sigma Aldrich) and sonicated for 5 minutes (Branson Sonicator). The concentrated emulsion was allowed to clear overnight. ApoC-III protein was solubilized in 4M guanidine hydrochloride and dialyzed against 10 mM NH_4_HCO_3_ buffer prior to the experiment. Protein concentration was assessed by bicinchoninic acid assay (Thermo Fisher Scientific). LPL-conditioned medium, generated by adenoviral transduction of human WT LPL in COS-7 cells was used as source of enzyme ^43^.

For each sample, concentrated lipid emulsion, apoC-III protein solution, and LPL-conditioned medium were combined to obtain the reaction mixture (final volume: 300 μl/sample).

The composition of the reaction mixture was: 3.4 mM Triolein (Sigma-Aldrich), approximatively 3.36μCi of ^3^H-triolein (PerkinElmer), 250 μM egg-phosphatidylcholine (Sigma-Aldrich), bovine serum albumin (0.75% mass/vol), heat inactivated human serum (5% vol/vol, used as source of ApoC-II), NaCl (0.15M), Tris HCl buffer pH=8 (0.05M), LPL-conditioned medium (100uL) and either WT, A23T or Q38K apoC-III protein solution (0-20μM in the final reaction mixture).

Reaction mixtures were prepared on ice and subsequently incubated at 37 °C for 1h. At the end of the incubation the products of reaction were extracted by adding 3.25 ml of methanol:chloroform:heptane solvent (1.41:1.25:1.00) and 1.05 ml of pH 10.0 Buffer Potassium Carbonate, Potassium Tetraborate, Potassium Hydroxide, Disodium EDTA Dihydrate (Thermo Fisher Scientific). Mixtures were centrifuged to allow the separation of the two liquid phases. 0.5mL of the upper phase, containing hydrolyzed fatty acids was counted (Beckman scintillation counter). The relative amount of hydrolysis was calculated for each sample and expressed as fraction of maximal LPL activity (0 μM of apoC-III). The IC_50_ values were calculated from the [inhibitor] vs. activity curves using a nonlinear regression model (GraphPad Prism statistical software).

### Generation of WT and Q38K encoding adeno-associated virus

The AAV8-TBG-APOC3 plasmid was generated as previously described ^13^. In order to obtain the AAV8-TBG-APOC3-Q38K, the Q38K variant was introduced onto the AAV8-TBG vector backbone by ligation-independent Gibson cloning. Briefly, the existing AAV8-TBG-APOC3 plasmid ^13^ was digested with Acc65I-HF and SalI-HF (New England Biolabs, Ipswich, MA), the vector backbone was gel purified, and the Q38K carrying APOC3 variant was introduced as a synthetic gene fragment hAPOC3_Q38K (Integrated DNA Technologies, Coralville, IA) using the NEBuilder HiFi DNA Assembly Master Mix according to the manufacturer’s instructions (New England Biolabs, Ipswich, MA). The sequence of the insert was validated by Sanger sequencing. AAV production, purification, and titration from the cloned WT and Q38K *APOC3* vector plasmids were performed by the Vector Core of the University of Pennsylvania.

### AAV WT versus Q38K APOC3 expression studies in mice

9-13 week old male mice were administered with either 1.6×10^12^ genome contents (GC/)mouse of WT or Q38K APOC3-AAV via intraperitoneal injection. 6 male mice per experimental group were used for these studies. For the determination of steady state lipid and ApoC-III plasma levels, plasma was collected prior to AAV administration (time 0, baseline) and at indicated time points. Blood samples were collected via retro-orbital bleeding using heparin-coated glass tubes after a 4h fasting period. The procedure was conducted under isoflurane-induced anesthesia. Lipid and apoC-III levels were determined using an Axcel autoanalyzer (Alfa Wassermann).

### Oral fat tolerance test (OFTT)

OFTT was performed in APOC3 WT and Q38K AAV treated mice at 4 weeks after AAV administration. Mice were fasted overnight and gavaged with 20 μl of olive oil (Sigma-Aldrich) per gram of fasting body weight. Blood was collected immediately before and at different time-points after olive oil administration (0-7h). TG levels were determined using a microplate commercial colorimetric assay (Infinity Triglycerides reagent). For the determination of TRL-bound ApoC-III, equal amounts of plasma from mice from each experimental group were pooled to a final volume of 60uL. Plasma was transferred to an ultracentrifuge tube (Beckman Coulter), layered with 1.34 mL of KBr solution d=1.006 and spun at 14,500 RPM for 25 min at 24°C to allow TRL floatation. The upper fraction, containing TRLs was collected. Total protein was precipitated using 100% Trichloroacetic acid (TCA) at a ratio of 1:4 (vol/vol). The precipitated protein was rinsed twice by adding 200 μl of cold acetone. After acetone removal, protein was resuspended in sample buffer containing 2-mercaptoethanol and separated by gel electrophoresis (4-12% gel, NuPAGE, and MES cathodic and anodic buffer, Thermo Fisher Scientific). Separated proteins were transferred onto nitrocellulose membranes, saturated with 5% milk solution and then incubated overnight at 4°C with a rabbit polyclonal antibody against human apoC-III (33A-R1b, Academy Biomedical) at a dilution of 1:1000 (vol/vol). Membranes were incubated with a secondary anti-rabbit IgG HRP conjugate antibody (NA934, GE healthcare) at a dilution of 1:10,000 for 1h at room temperature and bands corresponding to human ApoC-III were revealed using Luminata Crescendo chemiluminescent reagent (Millipore).

### Hepatic gene expression

Livers from AAV-injected mice were immediately collected after euthanasia, snap frozen and stored at −80°C until analyzed. RNA was extracted from liver samples from each mouse using TRIzol reagent (Thermo Fisher Scientific) according to the manufacturer’s protocol. 2 μg of RNA was reversed transcribed using the High-Capacity cDNA Reverse Transcription kit (Thermo Fisher Scientific). The gene expression levels of human *APOC3*, murine *Gapdh* and murine *36b4* were determined by quantitative real-time PCR, using a QuantStudio 7 Real-Time PCR System. Human *APOC3* gene expression was normalized to the mean Ct of the murine *Gapdh* and *36b4* gene expression from the same sample.

The primer sequences used for these reactions were:

*APOC3*: 5’-ACTCCTTGTTGTTGCCCTC-3’ and 5’-CGTGCTTCATGTAACCCTG
*Gapdh*: 5’-TGTGTCCGTCGTGGATCTGA-3’and 5’-CCTGCTTCACCACCTTCTTGAT-3’
*36b4*: 5’-TCATCCAGCAGGTGTTTGACA-3’ and 5’-GGCACCGAGGCAACAGTT-3’

### Assessment of ApoC-III Binding to plasma lipoproteins

Purified proteins were solubilized in in 4M guanidine hydrochloride and dialyzed vs 10 mM NH_4_HCO_3_ buffer prior to the experiment to a concentration of approximately 0.2–0.5 mg/ml.

Solubilized proteins were iodinated with ^125^I using the iodine monochloride method ^44^. Protein was iodinated with 1 mCi of ^125^I (PerkinElmer), 300 μl of 1 M glycine, and 150 μl of 1.84 M NaCl/2.84 μM ICl solution. The mixture was vortexed and loaded onto a PG-10 desalting column (Amersham Biosciences). Iodinated proteins were eluted in NaCl/EDTA solution and dialyzed against PBS. ^125^I activity was assessed by gamma counting (Packard Bioscience, Cobra gamma counter) and the protein concentration was assessed by bicinchoninic acid assay (Thermo Fisher Scientific). Human TRLs were isolated from pooled human plasma by density-gradient ultracentrifugation at d<1.006 mg/mL. Isolated TRL were dialyzed against PBS, and TG content was determined using a Cobas C311 autoanalyzer (Roche). For the assessment of apoC-III binding to plasma lipoprotein (whole plasma), 1 μg of ^125^I-apoC-III was combined with either 200 μl of pooled donor plasma or isolated TRL (380 mg/dL TG) and incubated at 37 °C for 1h. The relative binding of WT and Q38K apoC-III to whole plasma lipoproteins was assessed by FPLC fractionation (Superose 6 Increase gel-filtration column, GE Healthcare). TG and cholesterol concentrations in each fraction were determined using a microplate colorimetric assay (Infinity Triglycerides and Infinity Cholesterol reagents). ^125^I-apoC-III activity was determined by gamma counting (Packard Bioscience, Cobra gamma counter). The relative activity in lipoprotein fractions versus free protein fractions was assessed by comparing the relative magnitude of the area under the peaks corresponding to different lipoprotein fractions.

### Statistical analysis

Statistical analysis was performed using GraphPad Prism statistical software. Statistical comparisons between two groups were performed using a two-tailed Student’s t-test or two-way ANOVA as appropriate. Comparisons between three experimental groups were performed using one-way ANOVA or two-way ANOVA as appropriate. Statistical significance was defined as p< 0.05 for all analyses, after correction for multiple testing if appropriate.

## Results

### ApoC-III binding to TG-rich substrates is sufficient to impair TG hydrolysis *in vivo*

In order to test whether the binding of apoC-III to a TG-rich, hydrolysable substrate was sufficient to inhibit LPL activity and therefore impair TG clearance *in vivo*, we pre-incubated a commercially available TG-rich emulsion (Intralipid) with either recombinant human apoC-III (rhapoC-III) protein or PBS buffer and administered these substrates to *apoc3* KO and WT mice. As expected, in absence of any apoC-III (injection of PBS pre-incubated Intralipid into *apoc3* KO mice), the substrate was readily cleared from the circulation, whereas in the presence of apoC-III (injection of rhapoC-III pre-incubated Intralipid into apoc3 WT mice) we observed a significant delay in TG clearance (**figure 1**). Notably, the administration of rhapoC-III pre-incubated Intralipid to *apoc3* KO mice was sufficient to impair TG clearance to the same degree as that observed in *apoc3* WT mice (**figure 1**). This observation indicates that apoC-III binding to the substrate is sufficient to impair TG hydrolysis.

**Figure 1.**
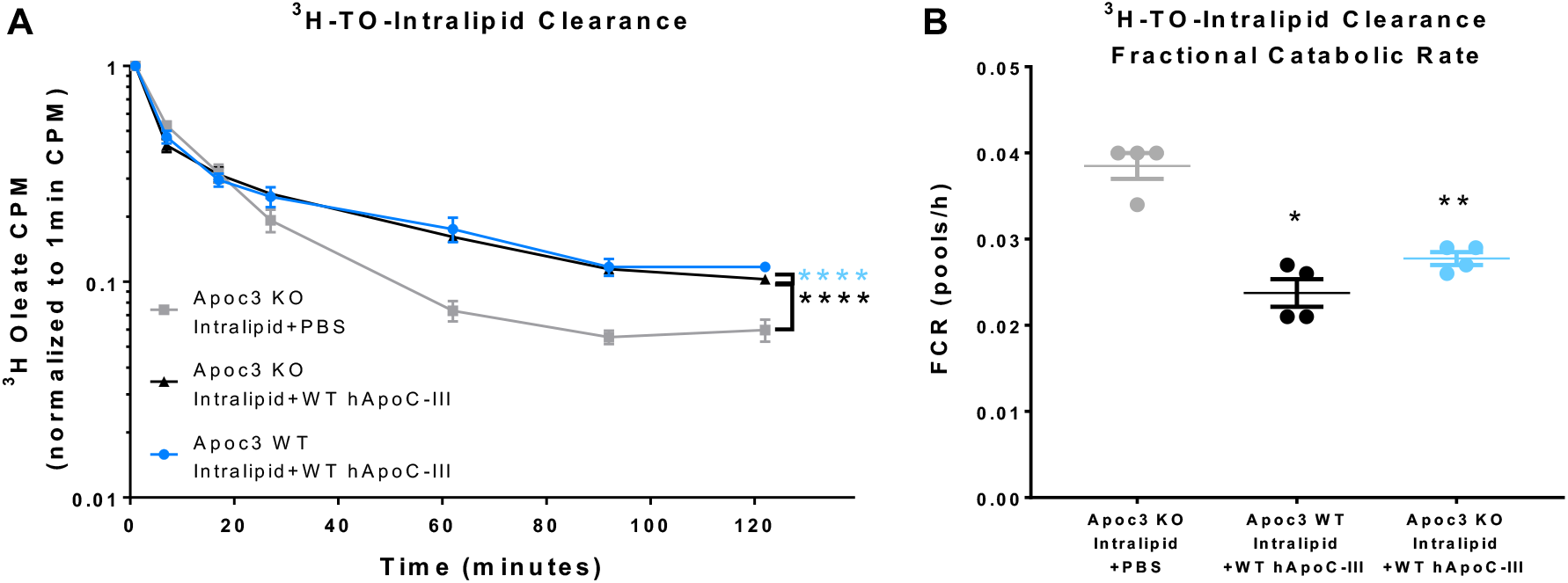
Assessment of the effect of ApoC-III binding on TG clearance *in vivo*. (**A**) Curves represent the ^3^H-TO-Intralipid clearance in apoc3 KO or apoc3 WT mice over the course of 122 min. Apoc3 KO mice were administered Intralipid that was pre-incubated with either PBS or WT rhapoC-III; apoc3 WT mice were administered Intralipid that was pre-incubated with WT rhapoC-III. ^3^H activity was expressed as normalized value, relative to plasma activity at 1 min. Data are shown as mean ± S.E.M per each experimental group (n=4 mice/group), y axis is represented in log scale. (A) ****P<0.0005, two-way ANOVA for the effect of the experimental treatment over time. Comparisons were tested vs apoc3 KO mice injected with PBS-pre-incubated Intralipid. (**B**) Fractional Catabolic Rate (pools/hour) for the curves shown in A. *P<0.05,.**P<0.01 one-way ANOVA vs Apoc3 KO mice injected with PBS-preincubated intralipid.

### Hydrogen exchange and mass spectrometry analysis indicates that apoC-III helical structure content and stability increase upon binding to lipoprotein sized substrates

It is apparent from the HX data that human apoC-III in the lipid-free state is unstructured because the amide hydrogens are unprotected and exchange at the theoretical rate expected for a dynamically disordered protein (**figure 2A)**. By this criterion, apoC-III is an intrinsically disordered protein which is consistent with prior CD studies showing very low α-helix content for this apolipoprotein^45, 46^. The HX data indicate that any helical structure present in the lipid-free apoC-III molecule must be highly unstable (**figure 2A)**. Interaction with a phospholipid (PL) - water interface enhances helix stability such that protection against HX is significantly increased, as shown in **figure 2A** for apoC-III associated with DMPC vesicles. Analysis of these HX time-courses for 9 peptides spanning the entire apoC-III amino acid sequence using procedures we have described before for the related apoA-I molecule^30^ gives the HX protection map depicted in **figure 2B**. It is apparent that the amide hydrogens in two central regions have Pf = 150 - 350 whereas the remainder of the amide hydrogens exhibit lower values. These Pf values indicate that the free energy of helix stabilization is about 2.5 - 3.0 kcal/mol which is lower than the value of 3 - 5 kcal/mol for the related apoA-I molecule in HDL particles^39^. The apoC-III α-helices are open ~ 0.5% of the time consistent with a highly dynamic and flexible structure for this protein. The Pf values tend to be lower when apoC-III is bound to more fluid lipid particles containing unsaturated PL acyl chains (palmitoyl-oleoyl phosphatidylcholine (POPC) SUV and POPC - triolein emulsion particles - data not shown) as compared to DMPC vesicles that contain saturated acyl chains. The terminal regions spanning residues 1 - 7 and 69 - 79 of the apoC-III molecule (**figure 2B**) have Pf = 1 indicating that they are dynamically disordered. Segments 8 - 14 and 46 - 47 have Pf in the 20 - 30 range which is consistent with either restricted coil structure or perhaps highly unstable α-helix. It is noteworthy that for peptides spanning the segments 15 - 41 and 48 - 65 the HX exchange parallels the intrinsic rate (**figure 2A**), showing that the complete segment exchanges by the same unfolding reaction (entire helix opening). The fact that high protection starts at amino acids 15 and 48 indicates that helices start at residues 11 and 44 since amides in the first turn of a α-helix are not protected against HX because they are not hydrogen bonded to the carbonyl groups of non-helical amino acids located 4 positions closer to the N-terminus of the protein.

**Figure 2.**
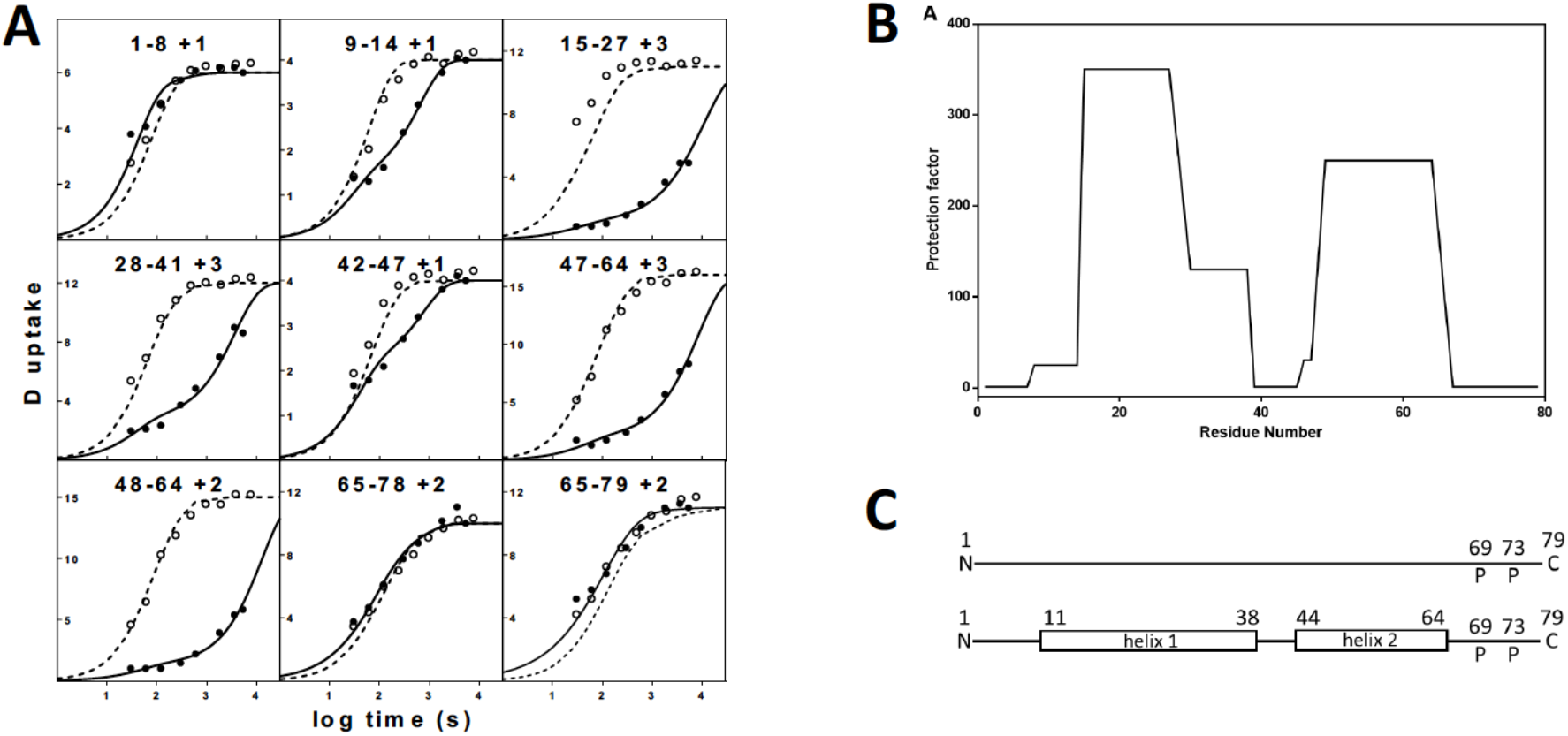
Comparison of HX kinetic profiles of human apoC-III in lipid-free and lipid-bound states. (**A**) HX kinetics (pD 5.8, 5°C, 20 - 40 μg/ml apoC-III) for 9 peptides (out of a total of about 50 unique peptides (many in multiple charge states) which provided internal consistency checks) covering the entire sequence of the apoC-III molecule are shown. In each panel (residue numbers and charge are indicated at the top) the observed HX kinetics (for amide hydrogens that undergo H-D exchange by EX2 kinetics - the rate of helix reclosing is faster than the rate of chemical exchange) of a lipid-free apoC-III fragment (○) and the same fragment associated with DMPC vesicles (apoC-III was incubated with the vesicles at room temperature for 1h at a 10/1 w/w lipid/protein ratio before the HX experiment was started) (●) are compared to the intrinsic rate for the peptide (dotted line). The intrinsic rate is the theoretically computed HX rate of the dynamically disordered random coil with Pf = 1. The time-courses for lipid-bound apoC-III are fitted to either stretched mono-exponential or bi-exponential rate equations. The former fit indicates a single cooperatively unfolding segment of secondary structure and the latter fit indicates a peptide that spans a helix terminus (7,16). The mass spectra for the experiments in this figure are shown in **figures S3** and **S4,** and the kinetic fitting parameters are listed in Table S2. (**B-C**) Summary of the HX-derived α-helix stabilities and secondary structure assignments for apoC-III in lipid-free and lipid-bound states. (**B**) Site-resolved stabilities for apoC-III associated with DMPC vesicles. The HX kinetic data in Fig. 1 and Table S1 were analyzed to obtain Pf values for each apoC-III peptide fragment. The Pf values are plotted for 9 peptide fragments (represented by a series of connected horizontal lines) that span the length of the apoC-III molecule. (**C**) The cylinders represent α-helices and the lines represent disordered structure. The positions of proline residues (P), whose presence leads to some perturbation of α-helix organization, are marked.

The locations of the α-helices in lipid-associated apoC-III are depicted in **figure 2B and C**. It is apparent that there are two 28 and 21 residue-long α-helices spanning amino acids 11 - 38 (helix 1) and 44 – 64 (helix 2), respectively. This information is the first report of the locations of α-helices in apoC-III molecules bound to lipoprotein-size lipid particles. This secondary structure is different to that of apoC-III associated with smaller SDS micelles where six ~ 10 - residue amphipathic α-helices are formed^47^. Comparison of the HX-derived secondary structures indicates that the α-helix content of apoC-III increases from zero to ~ 60% upon lipid binding (**figure 2C)**. Such an increase in helix content is broadly consistent with prior CD measurements^45, 46^ and with the fact that amphipathic α-helix formation provides much of the favorable free energy change that drives binding of exchangeable apolipoproteins to lipid-water interfaces^48^. Helix 2 spanning residues 44 - 64 is more hydrophobic and amphipathic (higher helical hydrophobic moment) than helix 1 that spans residues 11 - 38 and might be expected to have higher lipid affinity^49^. Consistent with this idea, after thrombin cleavage of apoC-III at position 40, the C-terminal fragment can form complexes with PL whereas the N-terminal fragment cannot^50^.

### The naturally occurring Q38K variant of apoC-III has increased LPL inhibitory activity *in vitro*

In our previous report, we showed that the effects of the LoF A23T apoC-III variant on plasma TG levels are consistent with these concepts ^13^; this variant displays impaired binding to plasma lipoproteins, accelerated clearance of the unbound protein from the circulation and as a result, reduced ability to inhibit TG clearance^13^. We hypothesized that conversely, a putative GOF variant in apoC-III could display increased binding to TRL and exert increased inhibition of LPL activity. To test our hypothesis, we decided to investigate the biochemical properties of Q38K, a naturally-occurring variant of apoC-III that has been associated with increased TG levels^22, 23^. We first investigated the inhibitory effects of WT, A23T and Q38K on LPL activity using a well-established *in vitro* assay^43^. Equal amounts of LPL were incubated with increasing concentration of ApoC-III and a TRL-like synthetic emulsion. Results indicated that as previously reported, A23T is less efficient that WT apoC-III in inhibiting LPL activity^13^, whereas Q38K displays significantly higher inhibition capacity (**figure 3A**). Thus, comparison of the half maximal inhibitory concentrations (IC_50_) for WT and variant apoC-III proteins indicates that a significantly higher concentration of A23T and a significantly lower concentration of Q38K are required to inhibit 50% of LPL activity (**figure 3B**).

**Figure 3.**
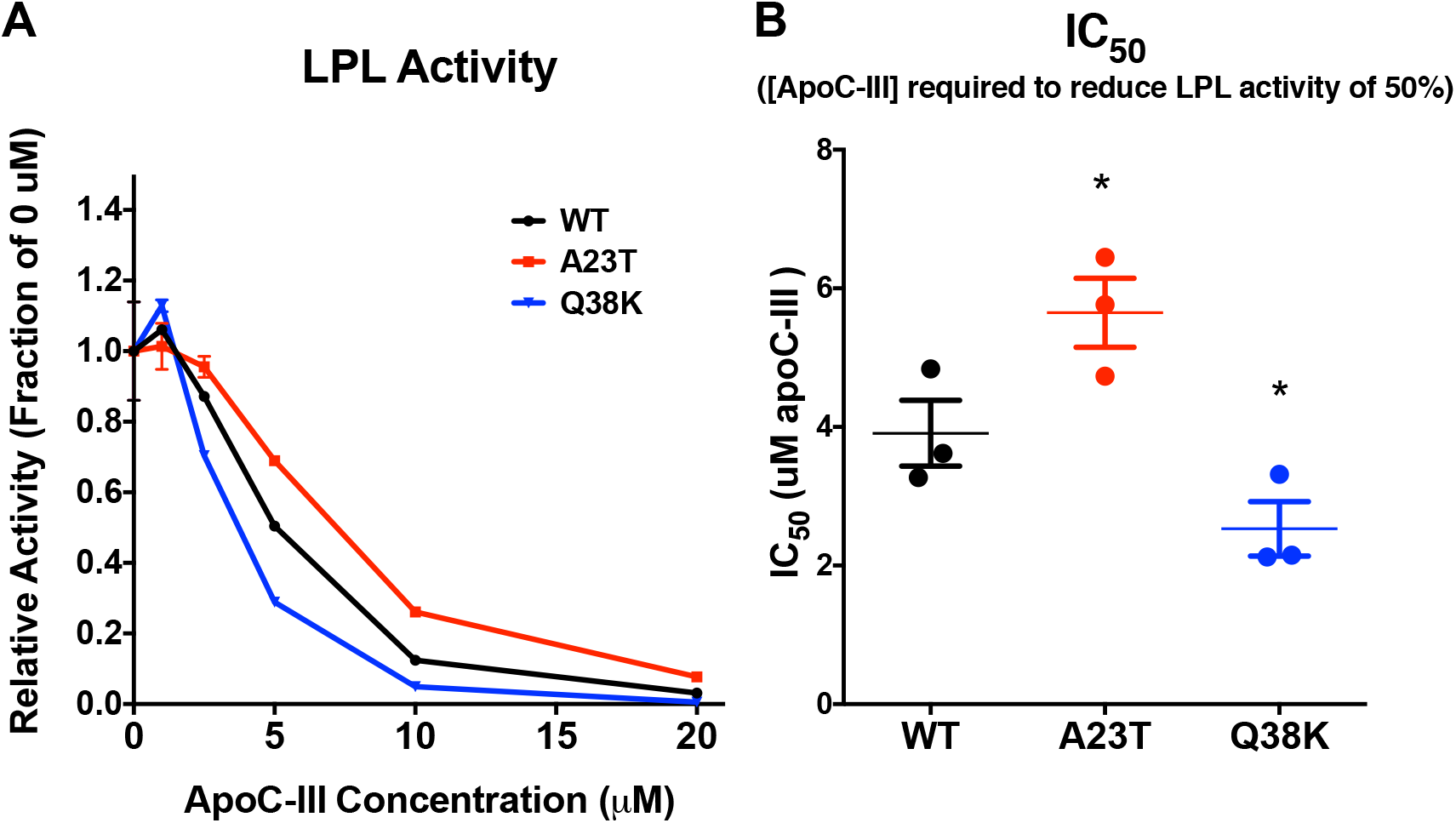
Comparison of *in vitro* LPL-inhibition capacity for WT, A23T and Q38K apoC-III. **(A)** Equal volumes of LPL conditioned medium were incubated with a TRL-like ^3^H-Triolein-containing emulsion and indicated concentrations of human apoC-III. Released ^3^H-fatty acids were separated by liquid-liquid extraction and quantified by scintillation counting. Points indicate the average value from two replicate samples from one representative experiment, Brackets indicate SEM. The experiment has been conducted three times. (**B)** IC_50_ for WT, A23T or Q38K human apoC-III. Bars indicate the average IC_50_ values from the three independent experiments. Brackets indicate SEM. *P<0.05, one-way ANOVA compared to WT apoC-III group.

### Expression of Q38K in *apoc3* KO mice induces hypertriglyceridemia and impaired TG lipolysis

To study the phenotypic effects of Q38K *in vivo*, we administered adeno-associated virus (AAV) vectors encoding either WT or Q38K human *APOC3* to *apoc3* KO mice. Both experimental groups displayed comparable levels of hepatic *APOC3* mRNA in response to the AAV treatment (**figure 4A**). Despite equal gene expression levels, mice expressing Q38K apoC-III displayed a significant increase in TG levels compared to mice expressing WT apoC-III (**figure 4B**). Mice expressing Q38K apoC-III showed a marginal but statistically significant decrease in the levels of total cholesterol and this decrease was due to a modest decrease in HDL-C levels (**figure S1A-C**). To investigate whether this hypertriglyceridemic effect of Q38K was due to impaired TG clearance, we exposed WT and Q38K AAV-injected mice to an oral fat tolerance test (OFTT). We observed that mice expressing Q38K apoC-III had significantly impaired postprandial TG clearance with an almost 2-fold increase in the area under the OFTT-TG curve (1.975 ± 0.03552, mean fold change ± SEM) compared to mice expressing WT apoC-III (**figure 4C and D**). Our previous data indicate that conversely, the expression of A23T in mice leads to accelerated postprandial TG clearance and to a significant reduction in AUC (e.g 0.4629 ± 0.04016 mean fold change ± SEM when expressed in WT C57BL/6 mice)^13^.

**Figure 4.**
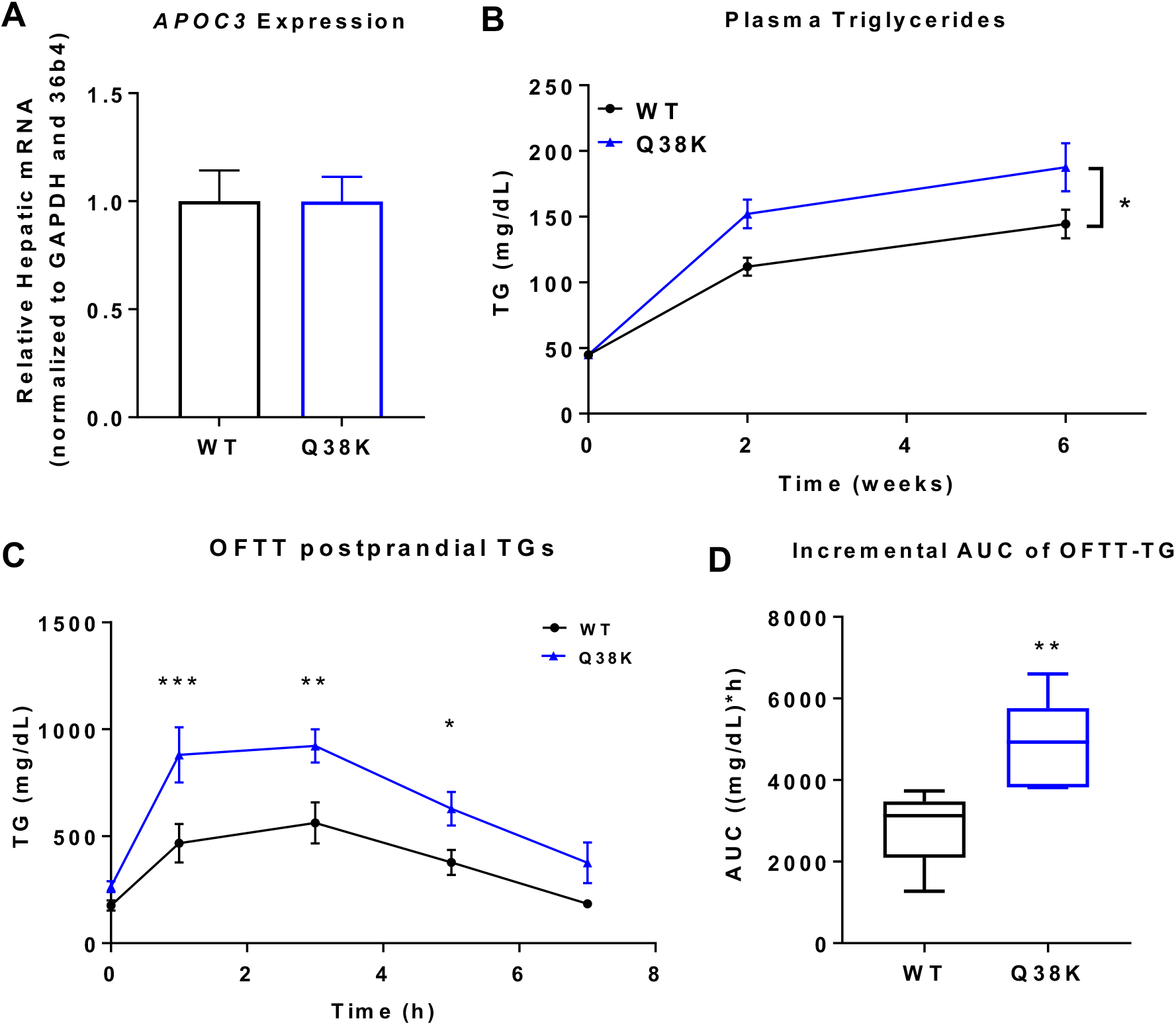
Comparison of the hypertriglyceridemic effects of WT and Q38K apoC-III *in vivo*. **(A**) 4h fasting plasma apoC-III in Apoc3 KO mice injected with 1.6 × 10^12^ GC/mouse of indicated *APOC3* AAV. Levels were measured 6 weeks after AAV administration. (**B**) Change in plasma TG levels in response to *APOC3* AAV administration in mice from A. (**C**) Postprandial TG clearance as determined by Oral Fat Tolerance test mice from A and B. (**D)** Incremental area under the curve (AUC) shown in C. (A-C) Data are expressed as mean ± S.E.M per each experimental group, (D) Boxes represent the 25th to 75th percentile range; the middle line represents the median. Whiskers extend to the minimum and maximum values within the group. The experiments (A-D) have been replicated on a separate cohort of mice, using a lower dose of AAV. (**B)** two-way ANOVA for the effect of the experimental treatment over time **(C**) *P<0.05, **P<0.01, ***P<0.001, two-way ANOVA with matching by time-point (**D**) **P<0.01, Student’s unpaired two-sided t-test.

### Q38K displays increased binding to apoB-containing lipoproteins *in vitro* and *in vivo*

We hypothesized that, despite similar gene expression levels, the Q38K variant may display increased binding to plasma lipoproteins and reduced removal from the circulation. To test this possibility, we measured apoC-III protein levels in mice from the experiment described in **figure 4.** ApoC-III protein levels were comparable in mice receiving WT or Q38K AAV (**figure 5A)**. These data indicate that in contrast to what was observed for A23T^13^, the Q38K variant does not impact apoC-III clearance from the circulation and the total levels of plasma apoC-III (**figure 5A**).

**Figure 5.**
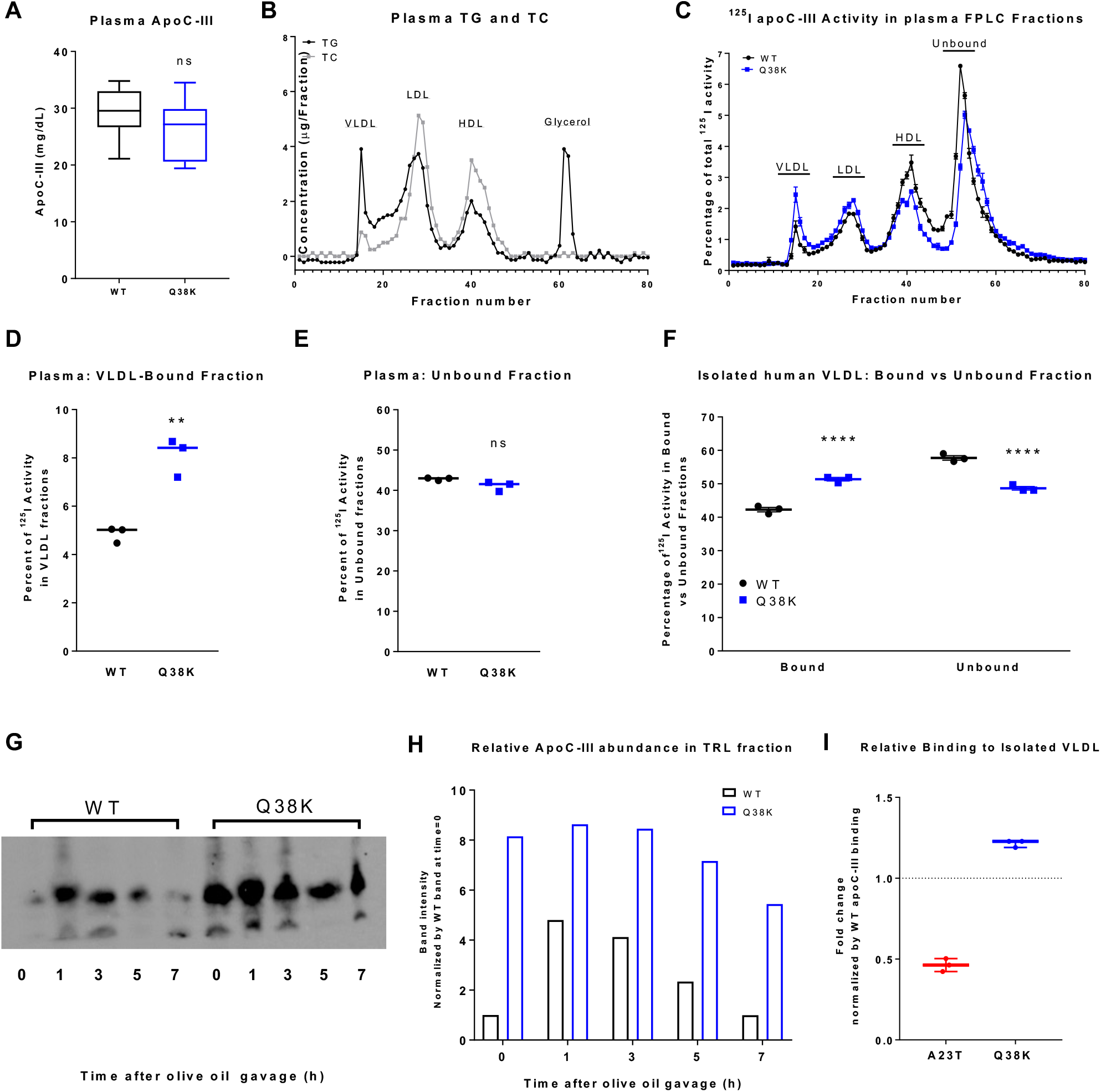
Assessment of the lipoprotein-binding capacity of WT vs Q38K ApoC-III. **(A**) Box plot of circulating apoC-III levels in 4h fasted mice from figure 4. (**B**) Cholesterol and Triglyceride content of FPLC-fractionated control human plasma used to assess apoC-III lipoprotein binding. (**C**) Relative binding of WT and Q38K apoC-III variants to lipoprotein fractions. 200 uL of human plasma was incubated with 1 μg of ^125^I-labeled WT or Q38K ApoC-III protein and fractionated using FPLC. (**D-E**). Relative percent of ^125^I activity in VLDL (D) and unbound fractions (E) from the curve in shown in (C). The relative percentage was estimated as area under the indicated peaks. Points represent the average of three technical replicates from one representative experiment. The experiment has been replicated three times. (**F**) Relative binding of WT and Q38K apoC-III variants to isolated human VLDL. A. 150 uL of isolated VLDL was incubated with 1 μg of ^125^I-labeled WT or Q38K ApoC-III protein and fractionated using FPLC. The relative percent of ^125^I activity in VLDL and unbound fractions was calculated as area under the relative peaks. Points represent the average of three technical replicates from one representative experiment. The experiment has been replicated twice (**G**) Immunoblot of apoC-III in isolated TRL from the OFTT described in **figure 4C and D**. Per each experimental group, samples collected at 0, 1, 3 and 7h after administration of olive oil were pooled and TRL were isolated by density gradient floatation (d<1.006g/mL). The supernatant was collected and blotted against human ApoC-III. The blot has been cropped to highlight bands corresponding to ApoC-III protein. Uncropped blot is shown in **figure S4**. (**H**) Densitometric assessment of the intensity of bands shown in the immunoblot in (G). Band densities were normalized by total loaded protein (intensity of Ponceau Red staining, **figure S4**). Results are expressed as intensity fold-change, relative to the first band (WT group at time 0). (**I**) Relative VLDL-binding capacity for Q38K and A23T. Q38K (panel I) and previously published A23T binding data^13^ were normalized by the binding capacity of their respective WT, control group and graphed side-by-side. (**B-I**) Data are shown as mean ± S.E.M, (**A**) Boxes represent the 25th to 75th percentile range; the middle line represents the median. Whiskers extend to min and max values within the group. **P<0.01, ****P<0.0001, Unpaired T-test.

To further understand this effect, we evaluated the ability of this variant to bind to lipoproteins by incubating human plasma with 1 μg of ^125^I-labeled WT or Q38K apoC-III protein and fractionating it using FPLC. The total cholesterol and TG content of separated FPLC fractions is shown in **figure 5B**. We observed that consistent with our *in vivo* data, there was no difference in the total amount of unbound protein (**figure 5C and E**). Interestingly however, Q38K showed increased affinity for apoB-containing lipoproteins, VLDL and LDL but decreased binding to HDL (**figure 5C, D and figure S2**).

In a complementary approach, we tested the binding of ^125^I-labeled WT or Q38K ApoC-III to isolated human TRL. Again, we observed that Q38K displayed increased TRL binding (**figure 5F**). To assess whether Q38K had higher binding capacity to endogenous TRL *in vivo*, we determined the relative amount of apoC-III in postprandial TRL fractions from *apoc3* KO mice expressing either WT or Q38K apoC-III. We observed that in both experimental groups, the total amount of TRL-bound apoC-III proportionally increased with the TG concentration in the sample and reached a peak in samples collected 1h after administration of olive oil (**figure 5G and H**).

Interestingly, the amount of TRL-bound apoC-III was higher in TRL isolated from mice injected with Q38K compared to WT APOC3 AAV at all time points (**figure 5G and H**). To allow a side-by-side comparison of the abilities of the Q38K and A23T variants to bind to TRL, we normalized re-plotted Q38K (panel I) and previously published A23T binding data^13^ by the binding capacity of their respective WT control group and graphed them side-by-side. It is apparent that the Q38K and A23T substitutions have opposite effects on apoC-III TRL-binding capacity (**figure 5I)**.

### The A23T and Q38K amino acid substitutions alter the hydrophilic-lipophilic properties of the amphipathic α-helices and ultimately affect the ability of apoC-III to bind lipoproteins

Equivalent experiments to those summarized in **figure 2** for WT apoC-III were performed for the loss-of-function variant A23T and the putative gain-of-function variants Q38K. The locations of the two α-helical segments in these mutated apoC-III molecules were not significantly different from those of the WT protein (data not shown). The results presented in **figure 6A** demonstrate that neither the A23T mutation nor the Q38K mutation significantly affects the stability of helix 1 which spans residues 11-38. The stability of helix 2 which spans residues 44-64 is also unaffected by these point mutations (data not shown). It follows that the altered lipoprotein-binding and LPL-inhibitory capabilities of these apoC-III variants are not a consequence of different overall α-helical structure but rather of altered interactions of the amphipathic α-helices with a PL-water interface. The helical wheel diagrams shown in **Figure 6B** and **C** describe the impact of the Q38K and A23T amino acid substitutions on the corresponding protein regions of WT apoC-III. The helical wheels depicted in **figure 6C** show that the A23T mutation leads to placement of a polar T residue in the nonpolar face of helix 1 that spans residues 11 - 38. The energy cost of inserting the hydroxyl group of the T sidechain into a hydrophobic lipid milieu leads to a reduced lipid-binding ability^13, 19^. In contrast, the Q38K point mutation leads to positioning of a K residue near the polar/nonpolar boundary of the amphipathic α-helix (**figure 6C**). Such placement of a K residue enhances the lipid affinity of α-helices because of a “snorkeling” effect^49, 51^. The four methylene groups of the K sidechain can remain in the nonpolar lipid environment while the polar amino group extends (“snorkels”) to interact with the aqueous phase. Such a conformational effect is consistent with the enhanced lipoprotein binding and LPL-inhibitory effects of the Q38K variant.

**Figure 6.**
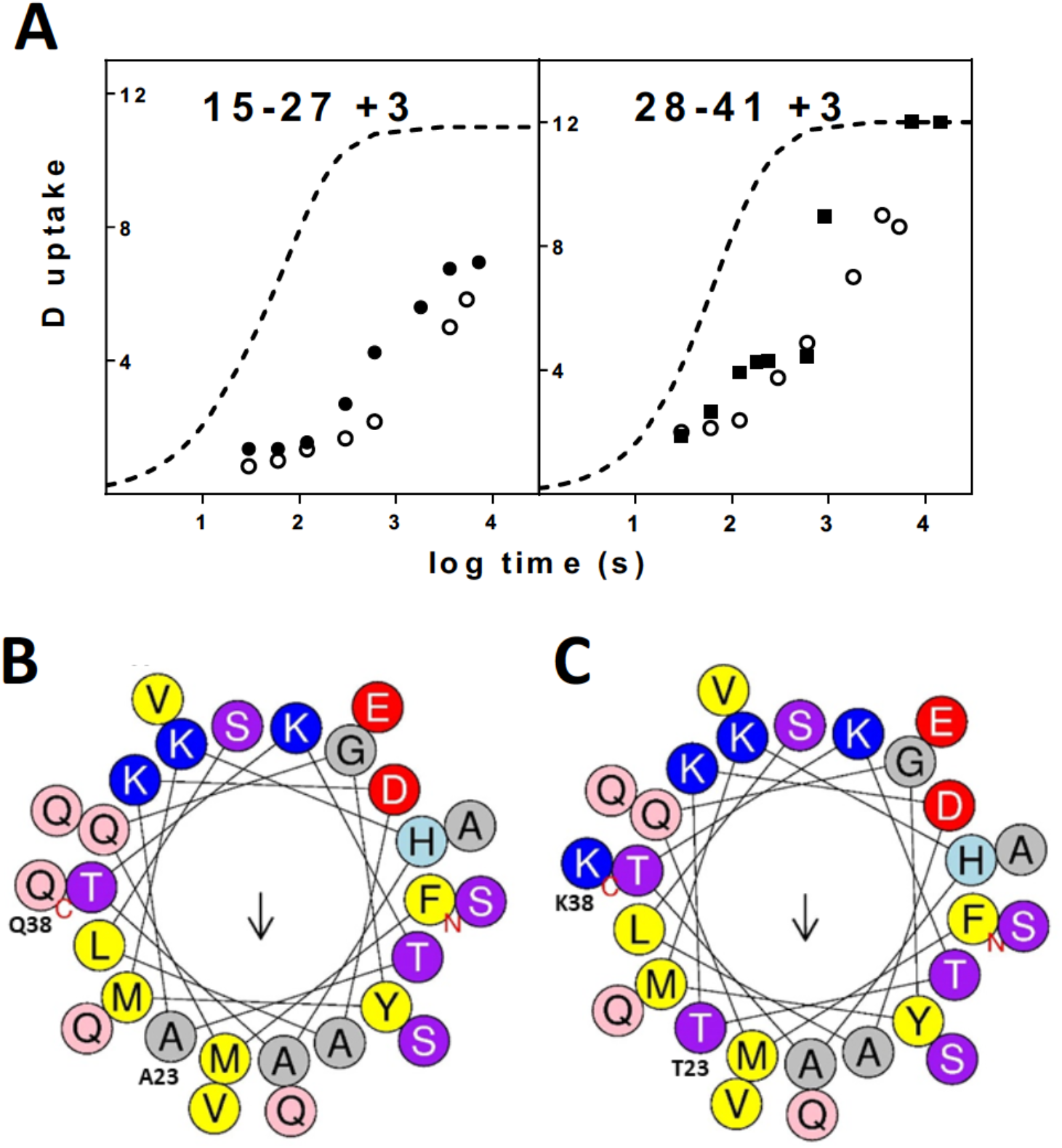
Comparison of HX kinetic profiles for peptides from WT apoC-III associated with DMPC vesicles and variants containing the mutations A23T and Q38K. **(A)** Left Panel. Peptide 15-27 that spans the A23T mutation site: (o) WT apoC-III, (●) A23T variant, dashed line is the intrinsic rate for the WT peptide. Right Panel. Equivalent data for peptide 28-41 that spans the Q38K mutation site: (o) WT apoC-III, (■) Q38K variant. (**B**) Helical wheel projection of residues 11-38 (helix 1) (FMQGYMKHATKTAKDALSSVQESQVAQQ) of WT human apoC-III with the N- and C-termini and the positions of A23 and Q38 indicated; the central vertical black arrow shows the direction of the helical hydrophobic moment and the nonpolar face of the amphipathic α-helix is at the bottom of the helical wheel. (**C**). The same segment of the apoC-III molecule with the T23 and K38 mutations included. It is apparent that the polar T residue is located in the nonpolar face of the amphipathic helix whereas the polar and charged K residue is located at the side of the helical wheel between the polar and nonpolar faces. The helical wheels were drawn with the HeliQuest webtool^56^.

## Discussion

Several studies have shown that circulating levels of apoC-III are inversely correlated with TG levels. This association is mediated by different and concurrent processes: apoC-III has been shown to increase VLDL secretion ^20, 21^, impair TG lipolysis within TRL^17, 18^ and reduce remnant uptake by the liver ^14–16^. The relative relevance of these pathways to the regulation of plasma TG is still unclear and likely affected by several other factors such as the studied experimental model, diet and comorbidity. In this study, we explored the mechanism of apoC-III interaction with TRL particles and its influence on LPL-mediated TG lipolysis.

LPL is the extracellular lipase responsible for the hydrolysis of plasma TG within VLDL and chylomicrons^52^. This process promotes the reduction of TG cargo within TRL and ultimately facilitates their catabolism. ApoC-III is known to inhibit LPL through different mechanisms. Earlier observations indicated that apoC-III may disturb the binding between TRL and glycosaminoglycans present on the cell surface thereby preventing efficient hydrolysis ^53^ whereas other sources indicate that apoC-III may compete with the LPL activator apoC-II for lipoprotein binding ^54^. In this setting, apoC-III may indeed actively displace apoC-II from the protein particle thus inducing impaired LPL activation^46, 54^. Both proposed mechanisms appear to share a common requirement for efficient binding of apoC-III to TRL. Our data support this observation and indicate that the binding of apoC-III to TRL-like particles is a crucial step in mediating LPL inhibition. In fact, the FCR of TG hydrolysis was significantly impaired in *apoc3* KO mice that received Intralipid pre-incubated with apoC-III, compared to control mice (*apoc3* KO mice that received Intralipid pre-incubated with PBS). Furthermore, this effect was apparently independent on the apoC-III content of other circulating lipoproteins. This conclusion is based on the observation that the substrate pre-incubated with apoC-III was cleared at similar rate in apoc3 KO and WT mice, regardless of the presence of native murine apoC-III.

In order to explore the mechanism of apoC-III binding to TRL, we examined the structural properties of WT human apoC-III in lipid-free and lipid-bound states using hydrogen-exchange mass spectrometry methods. Consistent with prior observations^45, 46^, we observed that the structure of apoC-III in the lipid-free state is disordered. Comparison of HX kinetic curves in lipid-free and lipid-bound states indicates that apoC-III forms amphipathic α-helix upon binding to a lipoprotein particle. Specifically, the lipid-bound protein contains 2 such helices that span amino acids 11-38 and 44-64.

In order to further investigate the relevance of this observations to apoC-III function, we investigated the ability of two naturally occurring variants to bind to TRL and impair LPL TG lipolysis. To this end, we intentionally selected two variants with opposite association with plasma TG: A23T and Q38K. As we previously reported, A23T is associated with reduced TG levels in humans and it has been shown to be a LoF variant of ApoC-III ^13, 19, 55^. The second variant Q38K was first identified in a family of Mexican descent and is associated with increased levels of TG^22^.

Our data show that the substitution of lysine for glutamine at position 38 also increases the LPL inhibitory capacity of apoC-III. This effect was observed in vitro, as demonstrated by increased LPL inhibition by Q38K compared to WT protein, and in vivo. Consistent with previous reports, the selective expression of Q38K *APOC3* in the liver of *apoc3* KO mice was associated with hypertriglyceridemia^23^ and we showed that this phenotype was associated with impaired postprandial TG clearance. These effects are consistent with our hypothesis that Q38K and A23T exert opposite effects on LPL-mediated hydrolysis of TG.

Our analysis also revealed that in mice Q38K apoC-III displays increased binding to TRL without changes in gene expression and total circulating levels of apoC-III protein. This effect seems to be explained by increased affinity for apoB-containing lipoproteins and reduced affinity for HDL particles. Our observations are consistent with prior reports from Sundaram and colleagues, who reported that in mice expressing Q38K apoC-III and injected with pluronic acid to inhibit hydrolysis of TG, apoC-III was mainly found in VLDL fractions and not HDL^23^. In this regard, the mechanisms of action of Q38K and A23T appear to be different. A23T displays reduced binding to all lipoprotein subclasses and as a consequence it is rapidly cleared by the kidney and removed from the circulation^13^. In contrast, the total levels of circulating Q38K do not seem to be affected and its accumulation on TRL particles appears to be explained by a shift in affinity towards apoB-containing lipoproteins.

In order to gain insights into the molecular basis of this phenotype, we applied hydrogen exchange-mass spectrometry to deduce how the mutations affect the structure and lipid binding properties of the protein. The LOF A23T variant and GOF Q38K variants are both located in the amphiphilic α-helix spanning residues 11-38 which contributes to the lipid binding ability of apoC-III. Neither substitution alters the helical structure of this protein segment but changes its hydrophilic/lipophilic properties. These modifications ultimately induce altered amphiphilicity of the α-helix and affect its lipid binding properties.

In summary, our study provides new mechanistic insights on the apoC-III binding to TRL and shows how this interaction is central to its LPL-inhibitory capabilities. Modifications of residues within helix 1 of apoC-III profoundly alter its lipid and lipoprotein-binding capacity. These data suggest that approaches aimed at disturbing the non-polar face of this specific protein region may offset the hypertriglyceridemic effects of apoC-III.

## Acknowledgments

The authors would like to thank David Nguyen for his outstanding technical expertise in the performance of the HX-MS experiments.

This work was supported in part by NIH grants R01HL133502 to D.J.R., by AHA Award 18POST34080184 to C.V. and by the NIH grant F31HL149162 to S.S.

