## Supplementary Material for "ApoC-III helical structure determines its ability to bind plasma lipoproteins and inhibit Lipoprotein Lipase-mediated triglyceride lipolysis"

### Supplemental Information

**Figure S1. Plasma lipid profile in apoc3 KO mice injected with either WT or Q38K APOC3 AAV.** Plots represent the plasma lipid levels for *apoc3* KO mice injected with 1.6 x $10^{12}$ GC/mouse of either WT or Q58K *APOC3* AAV, encoding for WT or Q38K human apoC-III, respectively. (**A**) Total Cholesterol, (**B)** HDL-Cholesterol (**C**) Non-HDL-Cholesterol plasma levels at indicated time-points after AAV administration. Data are expressed as mean ± S.E.M per each experimental group (n=6 mice/group). *P<0.05, **P<0.01, two-way ANOVA with matching by time-point.

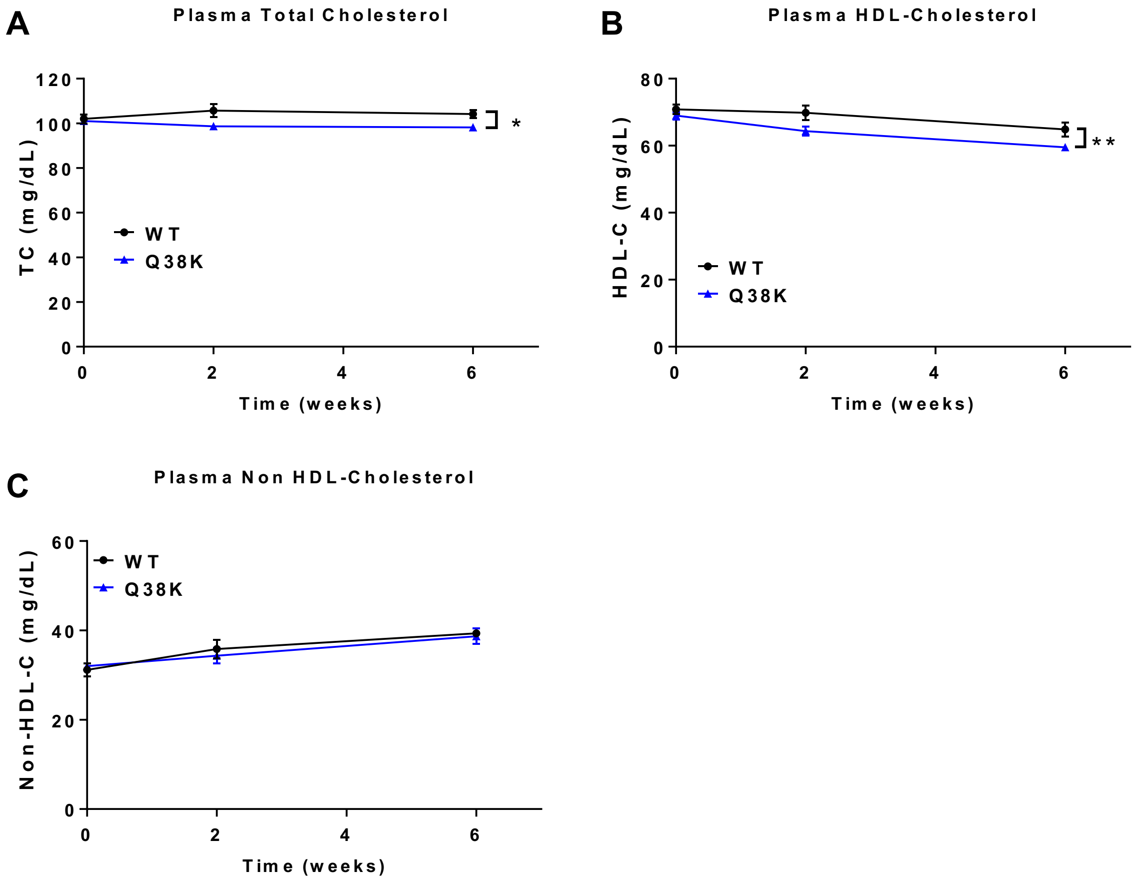

**Figure S2. Uncropped film and total protein staining for the blot shown in figure 5.** (**A**) Uncropped film of the image shown in **figure 5G**. (**B**) total protein staining (Ponceau Red staining) for the membrane used in (A). In order to permit the direct visualization of the reference molecular weight before and after immunoblotting, two different protein standard mixtures were loaded in the same well (first lane). The signal detected in (**A**) refers to an IgG-bound protein standard mixture, detectable by chemiluminescent reaction (MagicMark XP Western Protein Standard, Thermo Fisher). The signal shown in (**B**) refers to a pre-stained protein standard mixture (SeeBlue Plus2, Thermo Fisher).

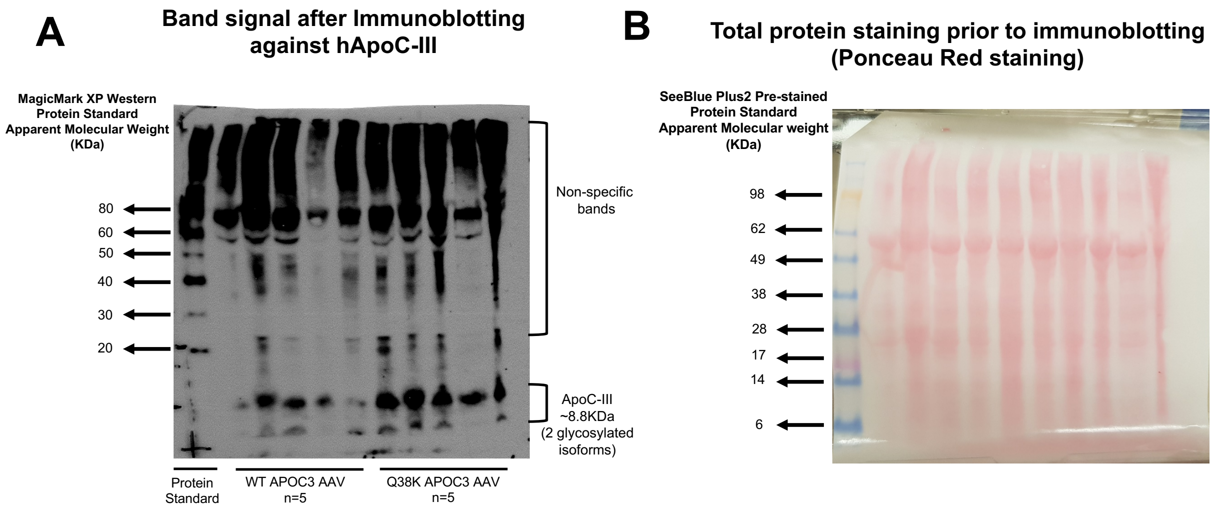

**Figure S3. Mass spectra of apoC-III peptides showing HX kinetics (pD 5.8, 5°C) in the lipid-free state**. The 9 peptides span the apoC-III amino acid sequence and the corresponding HX time-course are plotted in figure 2.

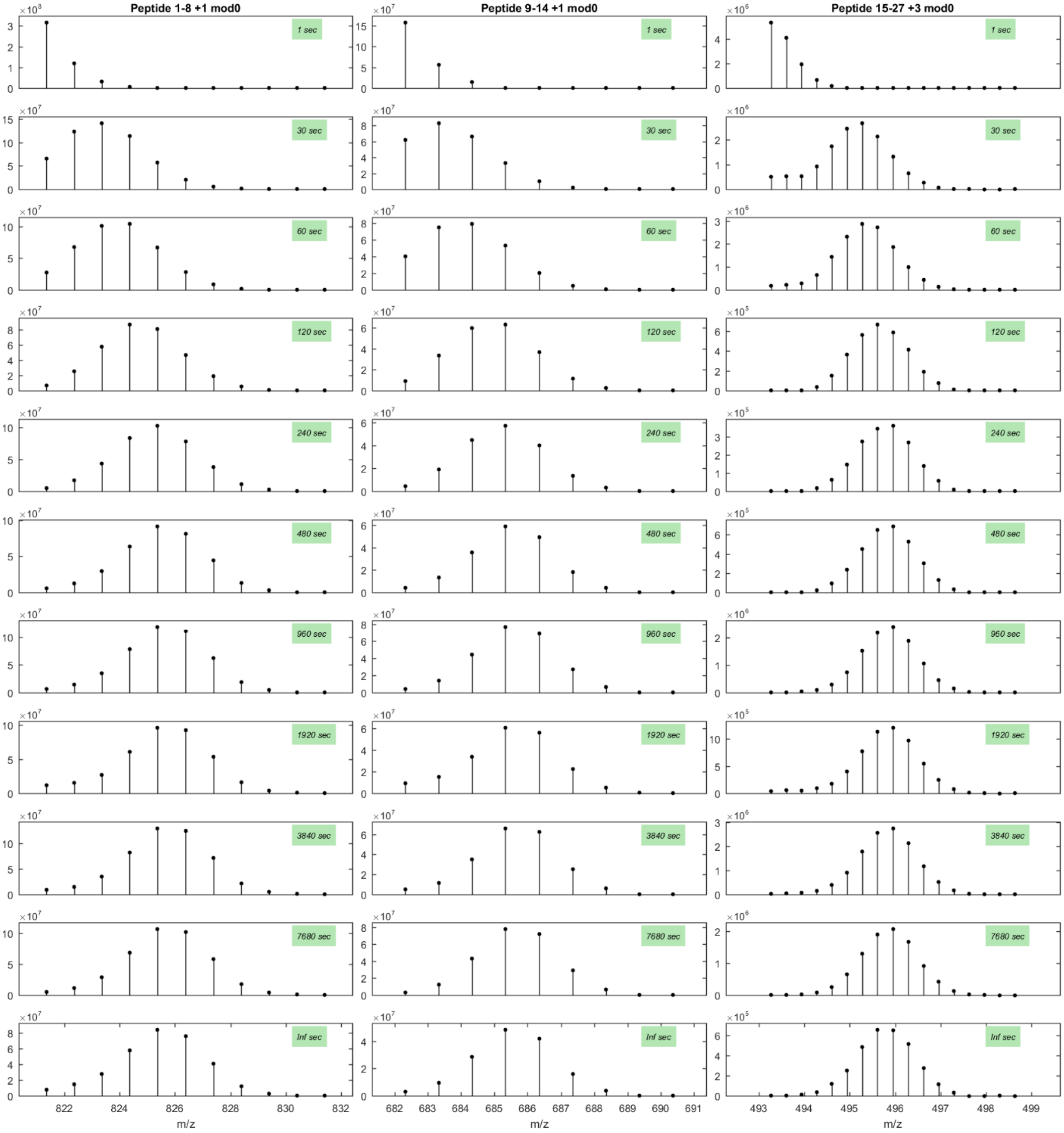

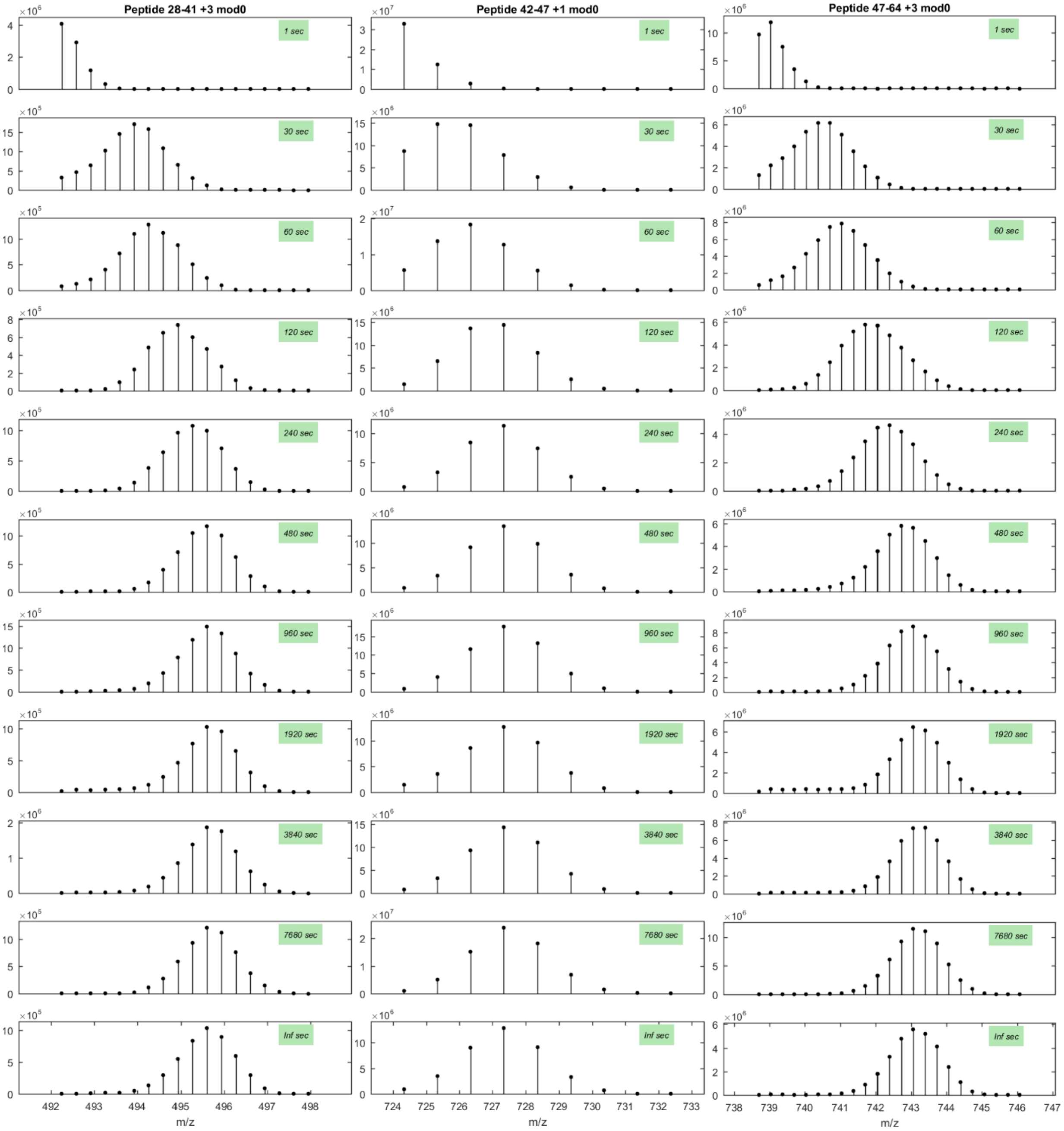

Peptide 28-41 +3 mod0
Peptide 42-47 +1 mod0
Peptide 47-64 +3 mod0
1 sec
30 sec
60 sec
120 sec
240 sec
480 sec
960 sec
1920 sec
3840 sec
7680 sec
Inf sec
m/z

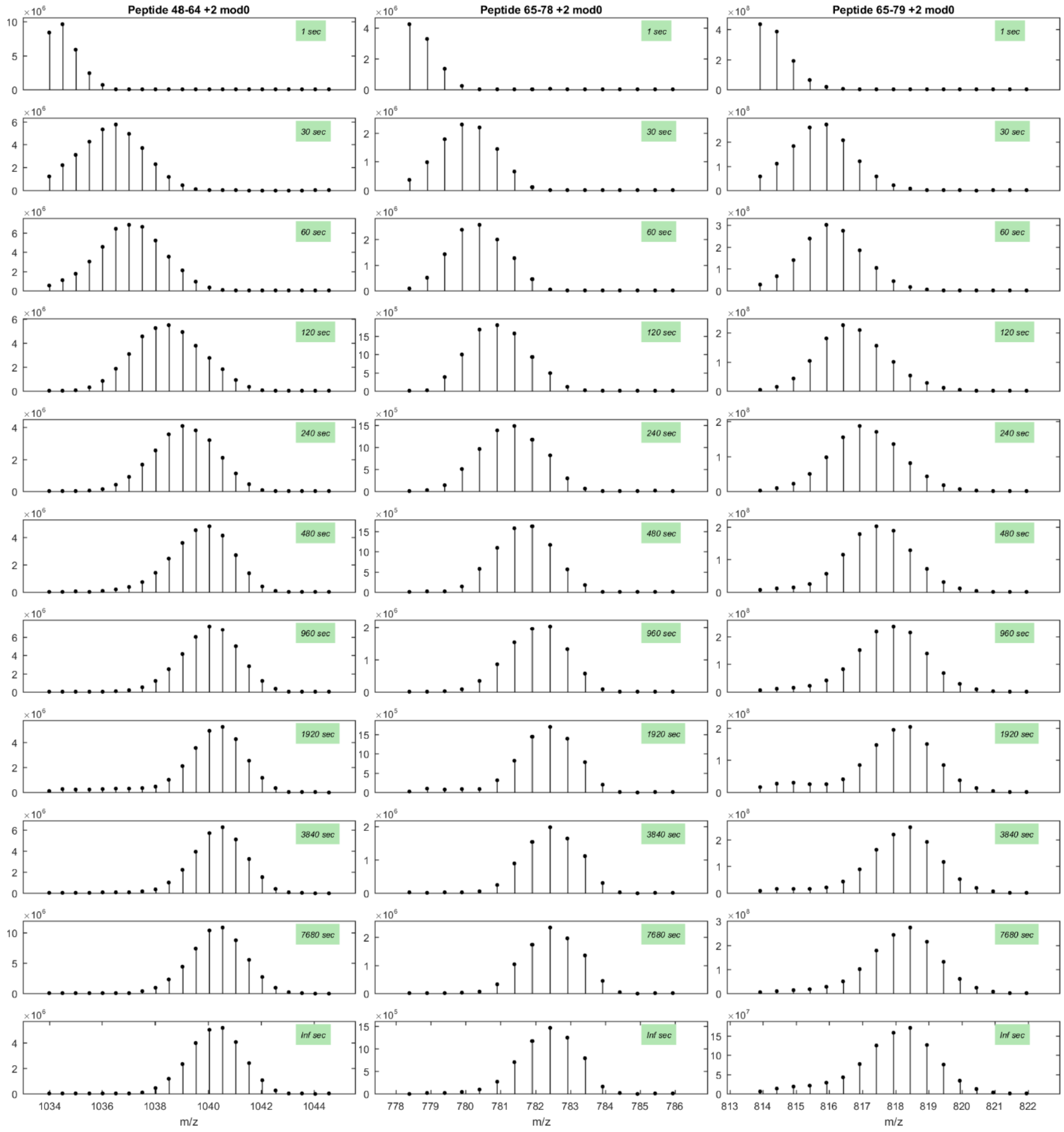
Peptide 48-64 +2 mod0
Peptide 65-78 +2 mod0
Peptide 65-79 +2 mod0
1 sec
30 sec
60 sec
120 sec
240 sec
480 sec
960 sec
1920 sec
3840 sec
7680 sec
Inf sec
m/z

**Figure S4. Mass spectra of apoC-III peptides showing HX kinetics (pD 5.8, 5°C) in the lipid-bound state**. The 9 peptides span the apoC-III amino acid sequence and the corresponding HX time-courses are plotted in figure 2. The mass spectra for certain peptide fragments are bimodal indicating that the intact protein exists as two conformations with fast and slow HX rates and therefore different degrees of D incorporation. To estimate the fractions of each population in the HX time-point samples, the mass spectra were fitted to a double Gaussian equation and integrated to obtain peak intensities. As we have discussed before for other apolipoprotein systems[1], helical segments located in the lipid-water interface are protected and exhibit slow HX kinetics; the reaction of this lighter peak follows EX2 kinetics (i.e. the rate of HX is much slower than the rate of helix refolding) so that the observed Pf value obtained by fitting the HX time-course (Table S1) is a measure of helix stability. The fast HX peak arises from protein segments that are desorbed from the lipid-water interface and are unprotected; the reaction of this heavier peak follows EX1 kinetics (i.e. the rate of HX is much faster than the rate of helix refolding) so that the Pf does not reflect helix stability.

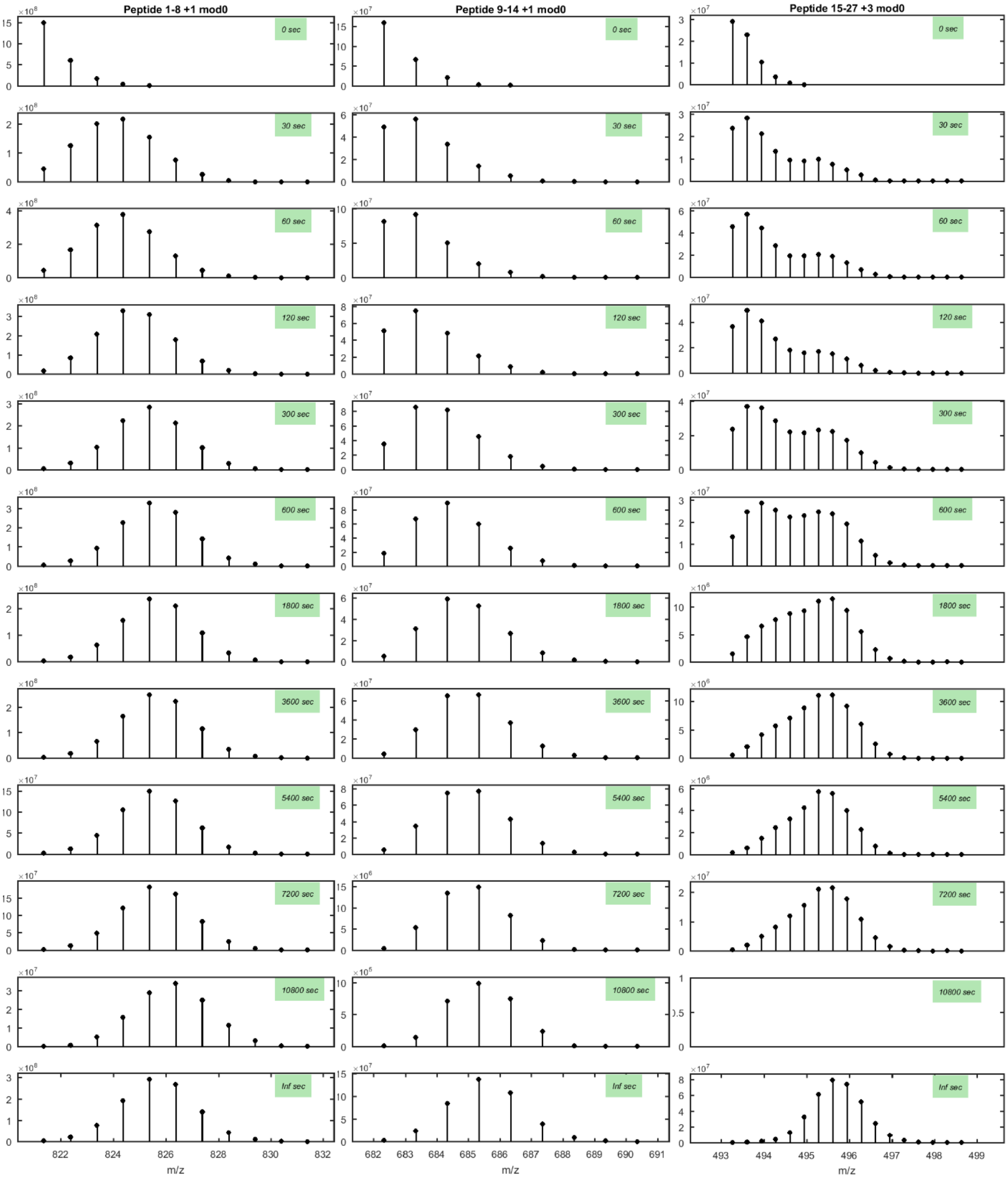
Peptide 1-8 +1 mod0
Peptide 9-14 +1 mod0
Peptide 15-27 +3 mod0
0 sec
30 sec
60 sec
120 sec
300 sec
600 sec
1800 sec
3600 sec
5400 sec
7200 sec
10800 sec
Inf sec
m/z
m/z
m/z

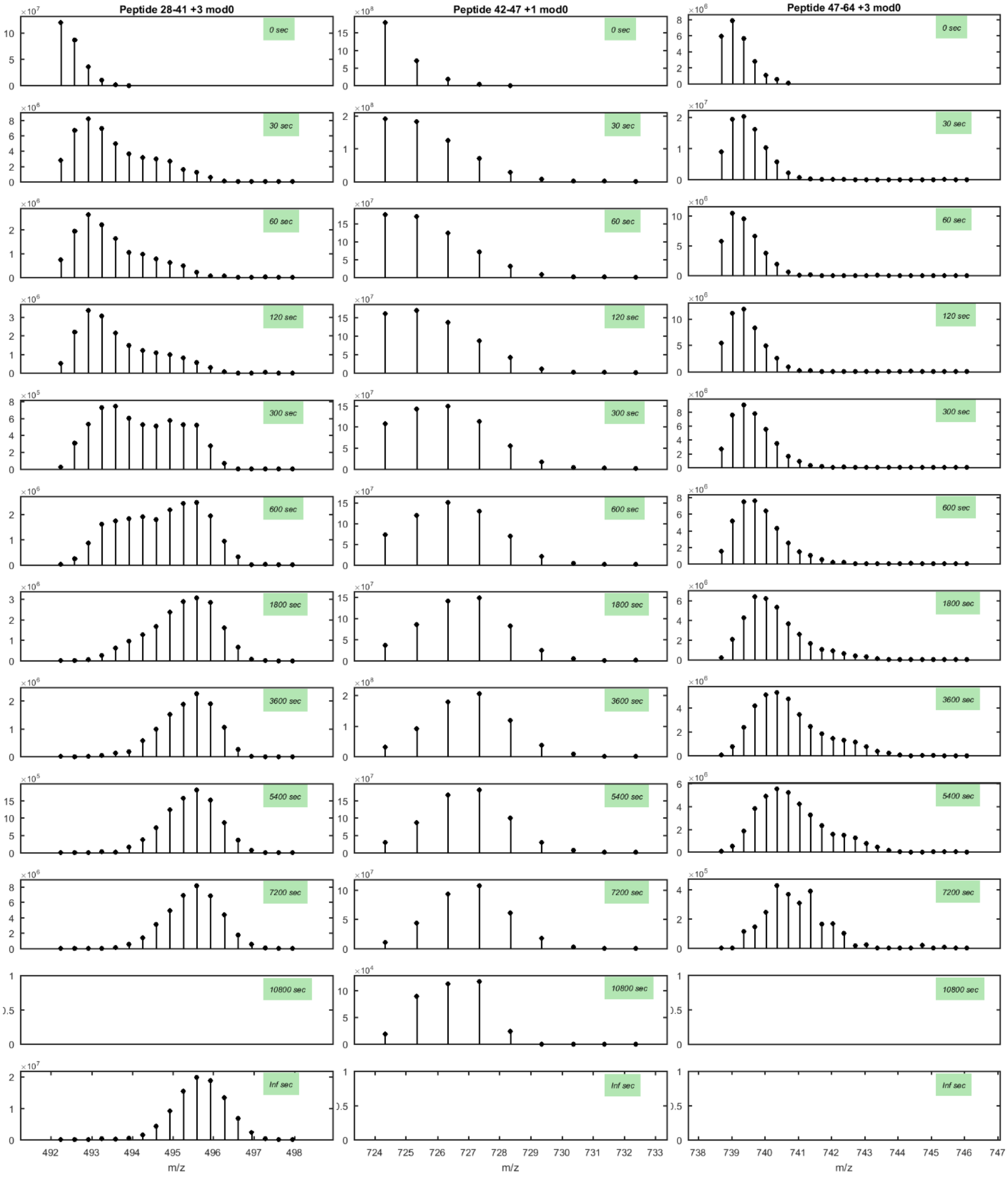
Peptide 28-41 +3 mod0
Peptide 42-47 +1 mod0
Peptide 47-64 +3 mod0
0 sec
30 sec
60 sec
120 sec
300 sec
600 sec
1800 sec
3600 sec
5400 sec
7200 sec
10800 sec
Inf sec
492 493 494 495 496 497 498
724 725 726 727 728 729 730 731 732 733
738 739 740 741 742 743 744 745 746 747
m/z
m/z
m/z

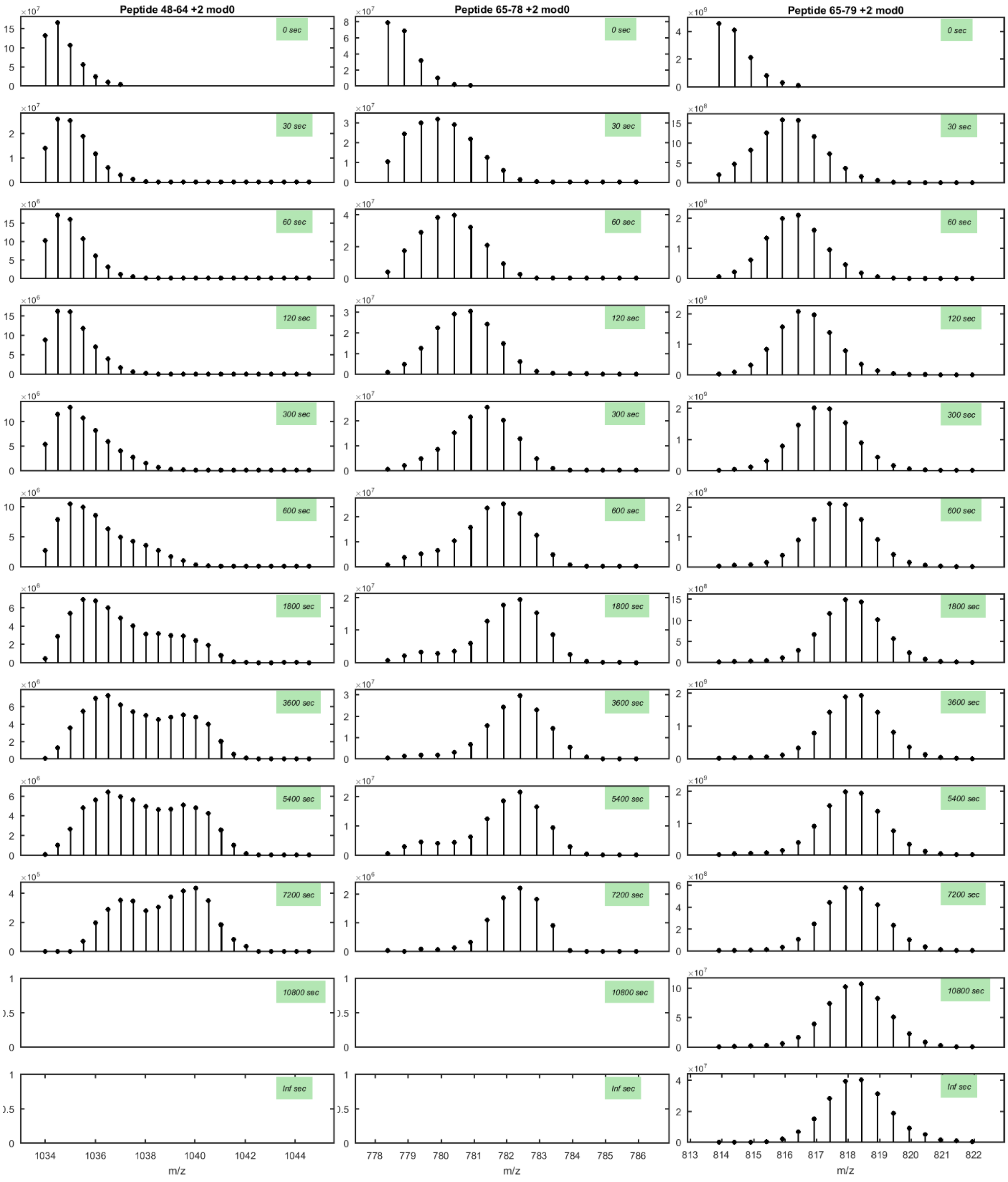
Peptide 48-64 +2 mod0
Peptide 65-78 +2 mod0
Peptide 65-79 +2 mod0
0 sec
30 sec
60 sec
120 sec
300 sec
600 sec
1800 sec
3600 sec
5400 sec
7200 sec
10800 sec
Inf sec
m/z

**Table S1. HX kinetics of apoC-III peptides associated with DMPC vesicles.** The HX time-courses obtained from the mass spectra of the 9 peptides included in figure S4 and plotted in figure 2 were fitted with one or two exponentials to derive the tabulated protection factors (Pf) and numbers of amides involved in each kinetic phase[1]. The fitting to the exponential equations was done using IGOR Pro (Wavemetrics Inc.).

| Peptide | Charge State | Stretch Factor (β) | Max D incorporated | # Unprotected Amides (Pf1) | # Protected Amides (Pf > 10) | Pf |
|---|---|---|---|---|---|---|
| 1-8 | 1 | 0.94 | 6 | 6 | 0 | |
| 9-14 | 1 | 0.94 | 4 | 1 | 3 | 25 |
| 15-27 | 3 | 0.78 | 11 | 1 | 10 | 350 |
| 28-41 | 3 | 0.89 | 12 | 3 | 9 | 100 |
| 42-47 | 1 | 0.92 | 4 | 2 | 2 | 20 |
| 47-64 | 3 | 0.84 | 16 | 2 | 14 | 200 |
| 48-64 | 2 | 0.88 | 15 | 1 | 14 | 280 |
| 65-78 | 2 | 0.75 | 10 | 10 | 0 | |
| 65-79 | 2 | 0.64 | 11 | 11 | 0 | |
